# SETD8 inhibition targets cancer cells with increased rates of ribosome biogenesis

**DOI:** 10.1101/2024.06.11.598464

**Authors:** Matilde Murga, Gema Lopez-Pernas, Robert Soliva, Elena Fueyo-Marcos, Corina Amor, Ignacio Faustino, Marina Serna, Alicia G. Serrano, Lucía Díaz, Sonia Martínez, Carmen Blanco-Aparicio, Marta Elena Antón, Brinton Seashore-Ludlow, Joaquín Pastor, Rozbeh Jafari, Miguel Lafarga, Oscar Llorca, Modesto Orozco, Oscar Fernández-Capetillo

## Abstract

SETD8 is a methyltransferase that is overexpressed in several cancers, which monomethylates H4K20 as well as other non-histone targets such as PCNA or p53. We here report novel SETD8 inhibitors, which were discovered while trying to identify chemicals that prevent 53BP1 foci formation, an event mediated by H4K20 methylation. Consistent with previous reports, SETD8 inhibitors induce p53 expression, although they are equally toxic for p53-deficient cells. Thermal stability proteomics revealed that the compounds had a particular impact on nucleoli, which was confirmed by fluorescent and electron microscopy. Similarly, *Setd8* deletion generated nucleolar stress and impaired ribosome biogenesis, confirming that this was an on-target effect of SETD8 inhibitors. Furthermore, a genome-wide CRISPR screen identified an enrichment of nucleolar factors among those modulating the toxicity of SETD8 inhibitors. Accordingly, the toxicity of SETD8 inhibitors correlated with MYC or mTOR activity, key regulators of ribosome biogenesis. Together, our study provides a new class of SETD8 inhibitors and a novel biomarker to identify tumors most likely to respond to this therapy.

## INTRODUCTION

Cancer genome sequencing efforts revealed that an important number of the driver mutations present in tumor cells occur on genes related to chromatin regulation (reviewed in [1]). These findings revitalized the efforts to develop drugs targeting epigenetic regulators (“epidrugs”), and today epigenetics is a very active area in the development of cancer therapies [2]. Initial evidences about the usefulness of targeting epigenetic regulators for cancer treatment came from the DNA methyltransferase (DNMT) inhibitor 5-Azacytidine (5-Aza), which already in 1976 showed effectiveness in clinical trials for acute myeloid leukemia (AML) [3]. Other successful examples include histone deacetylase (HDAC) inhibitors, approved for the treatment of cutaneous T cell lymphoma [4], or inhibitors of the Enhancer of Zeste 2 Polycomb Repressive Complex 2 Subunit (EZH2) histone methyltransferase, approved for rare sarcomas and follicular lymphoma.

SETD8 (aka KMT5A, PR-SET7 or SET8) is part of the SET (Su(var), Enhancer of zeste, Trithorax) family of histone methyltransferases, and the only one capable of catalyzing the methylation of histone H4 Lysine 20 (H4K20me1) [5]. This histone modification has been previously linked to a wide range of cellular functions such as chromatin compaction, transcription, mitosis or DNA repair (reviewed in [6]). In the context of genome integrity, SETD8 is essential for the recruitment of 53BP1 to DNA double strand breaks (DSBs), as this is mediated by binding of its TUDOR domain to methylated H4K20 [7]. Moreover, lysine methylation of PCNA by SETD8 regulates its stability and SETD8 inhibition reduces the levels of this factor that plays essential roles in DNA replication and repair [8]. Importantly, SETD8 also mediates the methylation of p53 (p53K382me1) marking it for degradation, and SETD8 inhibition has been proposed to kill cancer cells through p53-dependent apoptosis [9].

Evidence indicating that SETD8 could be an interesting target for cancer therapy has been building up in recent years. Initial work revealed overexpression of SETD8 in a wide range of cancers such as bladder cancer, non-small and small lung carcinoma, chronic myeloid leukemia, hepatocellular carcinoma and pancreatic cancer, where, as mentioned, its expression correlates with the levels of the DNA replication factor PCNA [8]. Other studies revealed a role for SETD8 in promoting the Epithelial to Mesenchymal Transition (EMT), suggesting that SETD8 inhibition could be particularly relevant for preventing cancer metastasis [10]. Arguably, the concept of targeting SETD8 in cancer therapy gained significant momentum with a study published in 2017, where chemical and genetic screens converged to identify SETD8 as a specific vulnerability in High-Risk Neuroblastoma [9]. Shortly thereafter, a similar approach revealed that SETDB8 was also a selective target in MYC-driven Medulloblastoma [11]. However, a mechanism that explains why SETD8 inhibition is particularly toxic for these tumors is currently missing. We here report the discovery of a new class of SETD8 inhibitors and provide mechanistic analyses that indicate that this therapy is particularly effective in tumors with high rates of ribosome biogenesis, such as those driven by the MYC oncogene.

## RESULTS

### An *in silico* screen identifies compounds inhibiting 53BP1 foci

Our original aim was to identify compounds that prevented the recruitment of the DNA repair factor 53BP1 to DSBs, an event that can be visualized as the accumulation of nuclear foci at sites of DNA damage [12]. To do so, we performed a virtual screen of around 1 million compounds (see Methods), using a set of available PDB structures of the TUDOR domain of 53BP1 bound to an H4K20me2 peptide [13], aiming to discover chemicals capable of occupying the same pocket and thereby compete with histone peptide binding. 25 selected compounds (**Table S1**) were then experimentally evaluated by assessing their effect at 10 μM on the formation of 53BP1 foci by High Throughput Microscopy (HTM), in human osteosarcoma U2OS cells exposed to 10 Gy of ionizing radiation (IR) (**Fig. 1A**). These experiments identified a single compound that prevented the formation of 53BP1 foci (**Fig. 1B,C**), which we named as compound 23 (C23). Subsequent modifications around the chemical scaffold of this compound identified additional compounds with increased potency in preventing 53BP1 foci formation such as compounds C110 or C111 (**Fig. S1**).

**Figure 1.**
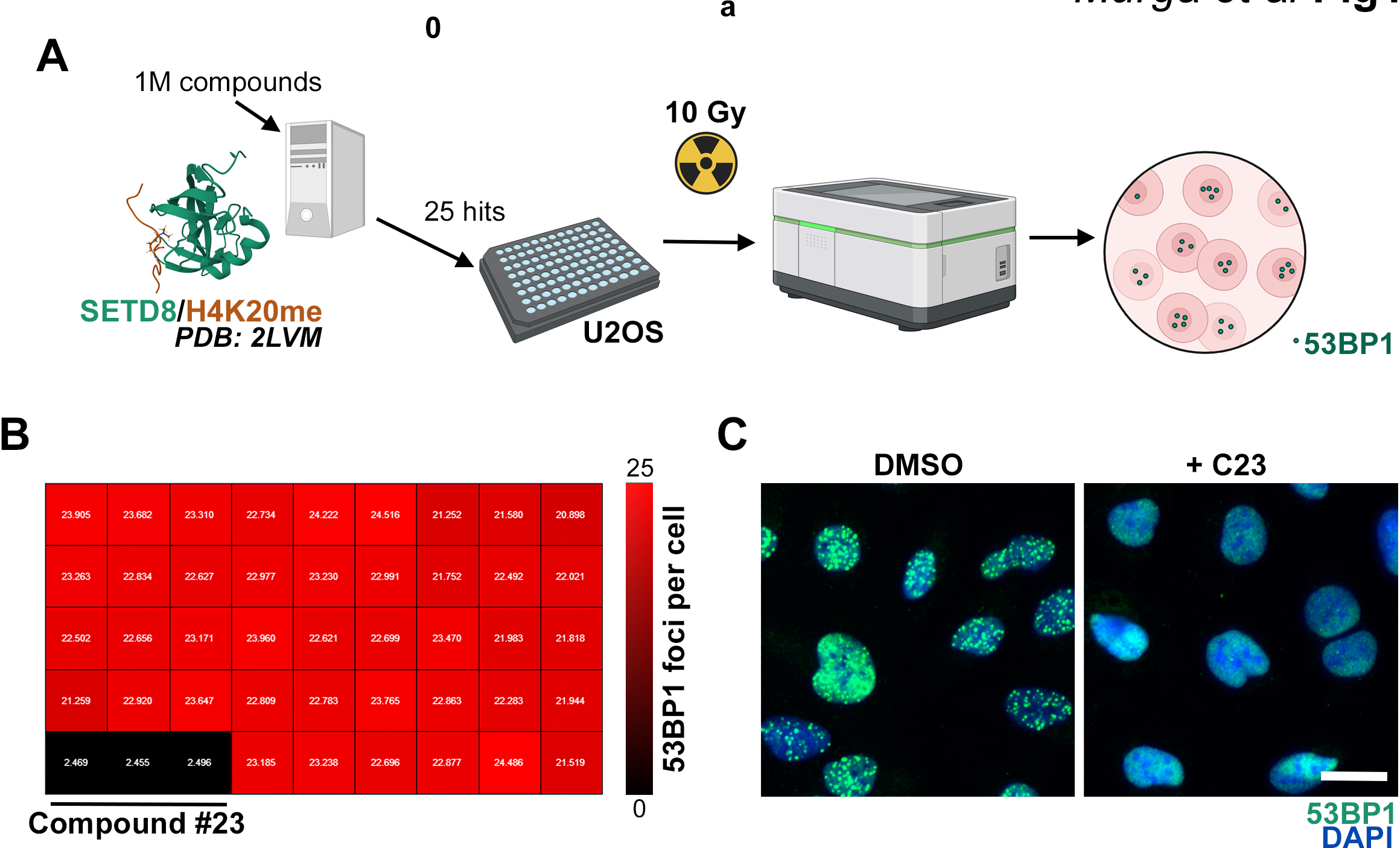
A virtual screen to identify compounds inhibiting 53BP1 foci. (**A**) Pipeline of the screening strategy. A virtual screen of around 1M compounds was performed to find chemicals mimicking the binding of a methylated H4K20 peptide bound to the TUDOR domain of 53BP1, using a published structure (PDB: 2LVM). 25 hits were next tested by HTM in U2OS cells exposed to 10Gy and evaluated for 53BP1 foci formation 45 min post-IR. (**B**) Heatmap (red-to-black) illustrating the number of 53BP1 foci per nucleus in each well of a plate from the screen, where each compound was tested in triplicate. Note that there was only a clear hit (C23) that abrogated 53BP1 formation. (**C**) Representative image from the experiment defined in (**B**), where 53BP1 foci (green) were analyzed by immunofluorescence. DAPI (blue) was used to stain DNA. C23 was used at 10 μM. Scale bar (white) indicates 5 μm.

### Identified compounds inhibit 53BP1-dependent functions

To determine to what extent C23 affected 53BP1 function, a series of additional experiments were conducted. First, the formation of 53BP1 foci depends both on the interaction of its TUDOR domain with methylated H4K20, as well as on ATM-dependent phosphorylation of targets such as histone H2AX [14]. Importantly, while C23 inhibited IR-induced 53BP1 foci, it did not affect the formation of foci for phosphorylated H2AX (γH2AX) in U2OS cells (**Fig. 2A**). Similar observations were made with additional compounds from the same chemical series (**Fig. S1B**). Consistent with immunofluorescence data, western blotting (WB) failed to detect any effect of C23 in IR-induced phosphorylation of CHK2 or KAP, two well-established ATM-dependent phosphorylation events (**Fig. 2B**). These results suggested that the compound was selectively affecting the TUDOR-H4K20me route for 53BP1 recruitment. Of note, C23 did not generate spontaneous DNA damage, as evidenced by the lack of KAP or CHK2 phosphorylation in non-irradiated U2OS samples treated with increasing concentrations of the drug (**Fig. 2B**).

**Figure 2.**
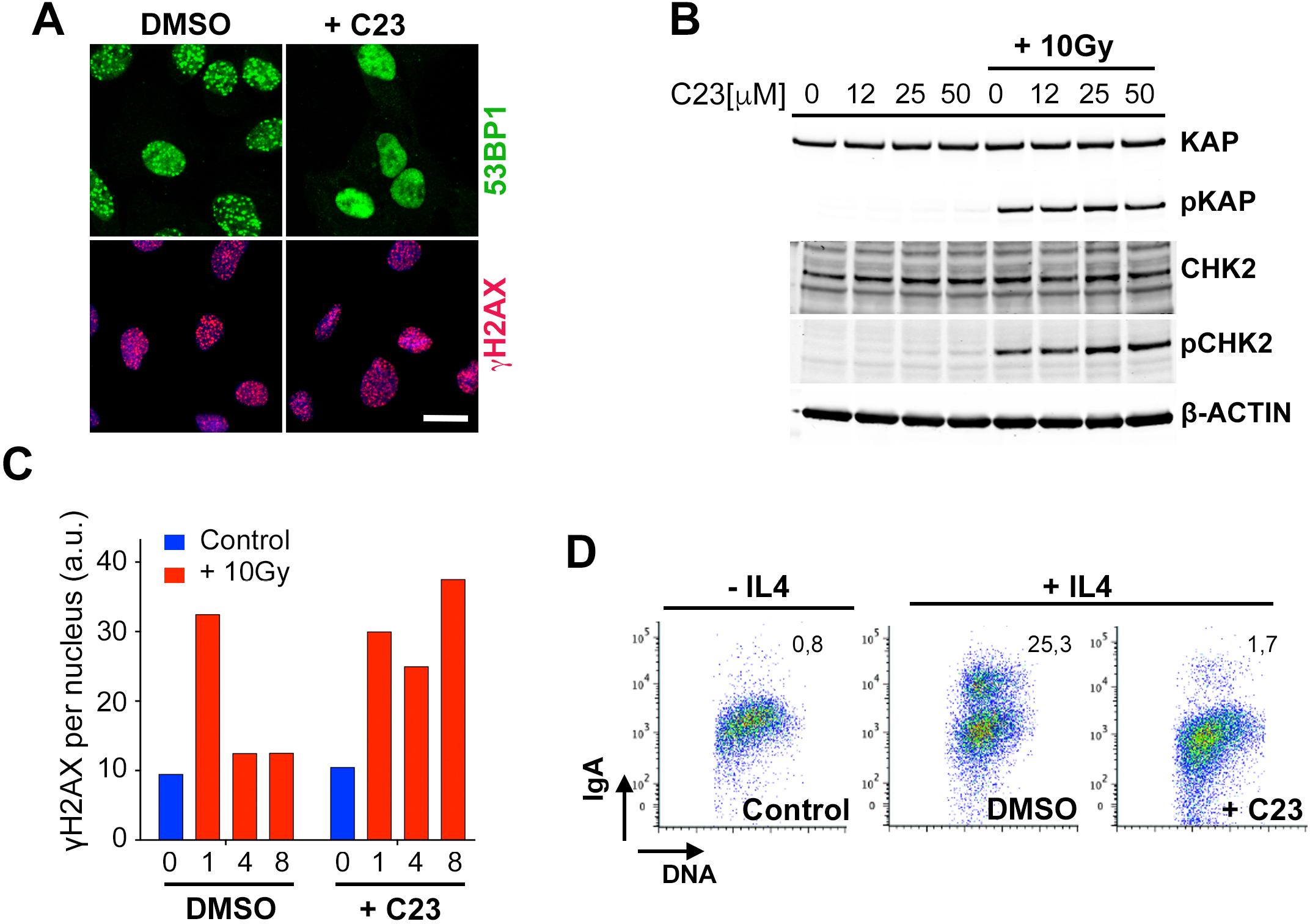
C23 impairs 53BP1 functions. (**A**) Immunofluorescence of 53BP1 (green) and γH2AX (red) in U2OS cells, treated or not with C23 (10 μM), 45 min after exposure to 10Gy of IR. Scale bar (white) indicates 5 μm. (**B**) WB of KAP, phosphorylated KAP (pKAP), CHK2 and phosphorylated CHK2 (pCHK2) in U2OS cells 45 min after exposure to 10 Gy of IR, in the presence or absence of the indicated concentrations of C23. Levels of β-ACTIN are shown as a loading control. (**C**) HTM-dependent quantification of the mean signal of γH2AX per nucleus, in U2OS cells at various times after being exposed to 10 Gy of IR, in the presence or absence of C23 (10 μM). (**D**) Flow cytometry in CH12 cells illustrating the effect of C23 (10 μM) in the efficacy of IL4-induced CSR (as measured by IgA expression). Numbers indicate the percentage of IgA+ cells.

Next, we evaluated if C23 could inhibit DNA repair. HTM-dependent quantification of nuclear γH2AX intensity in U2OS cells exposed to 10Gy of IR, showed that the drug delayed the clearance of the γH2AX signal, reflecting a generalized impairment of DSB repair (**Fig. 2C**). To focus on a DNA repair reaction that is specifically dependent on 53BP1, we analyzed class switch recombination (CSR), a B-cell specific reaction that involves the joining of broken DNA ends [15]. To do so, we used CH12 cells, a murine B cell lymphoma that undergoes efficient CSR from IgM to IgA in response to interleukin 4 (IL4) [16]. Remarkably, treatment with C23 led to a profound inhibition of CSR in CH12 cells (**Fig. 2D**). Together, these experiments indicate that C23 inhibits 53BP1 recruitment to DSBs and 53BP1-dependent repair activities.

### C23 is a SETD8 inhibitor

While conducting the DNA repair analyses mentioned above, we noticed that C23 also limited cell proliferation, which cannot be mediated by 53BP1 as this factor is dispensable for DNA replication and cell growth [17]. In contrast, C23 led to a dose-dependent reduction in DNA replication, as assessed by quantifying the incorporation of 5-Ethynyl-2′-deoxyuridine (EdU), which was equally observed in 53BP1^+/+^ and 53BP1^-/-^ mouse embryonic fibroblasts (MEF) (**Fig. S2A**). These results indicated that C23 was inhibiting 53BP1 recruitment to DSBs through an indirect effect.

The original virtual screen was designed in a way that, potentially, could identify compounds that compete with H4K20 binding in other targets besides 53BP1. In this context, we reasoned that the drug could be inhibiting SETD8, the methyltransferase responsible for H4K20 monomethylation, and thus with a binding site structurally similar to the one used for the virtual screen. Such a target could help to explain the effects of C23 on DNA replication, as SETD8 has been associated with DNA replication by the methylation of non-histone factors such as PCNA [8]. In support of this, an HA-tagged SETD8 (SETD8^HA^) co-localized with PCNA in U2OS cells (**Fig. 3A**). Moreover, the effect of C23 on DNA replication was equivalent to that observed upon treatment with UNC0379, a widely used SETD8 inhibitor [18] (**Fig. 3B**). Quantification of IR-induced 53BP1 foci in U2OS cells exposed to 10 Gy also showed a similar effect for UNC0379 and C23 (**Fig. 3C**). Besides DNA replication, another phenotype that has been associated to SETD8 deficiency is the accumulation of polyploid cells and cells arrested in G2, which arise from segregation problems during mitosis [19]. Consistent with this, flow cytometry analysis of DNA content showed that treatment with C23 or UCN0379 led to a progressive accumulation of G2 arrested and polyploid cells (**Fig. 3D**).

**Figure 3.**
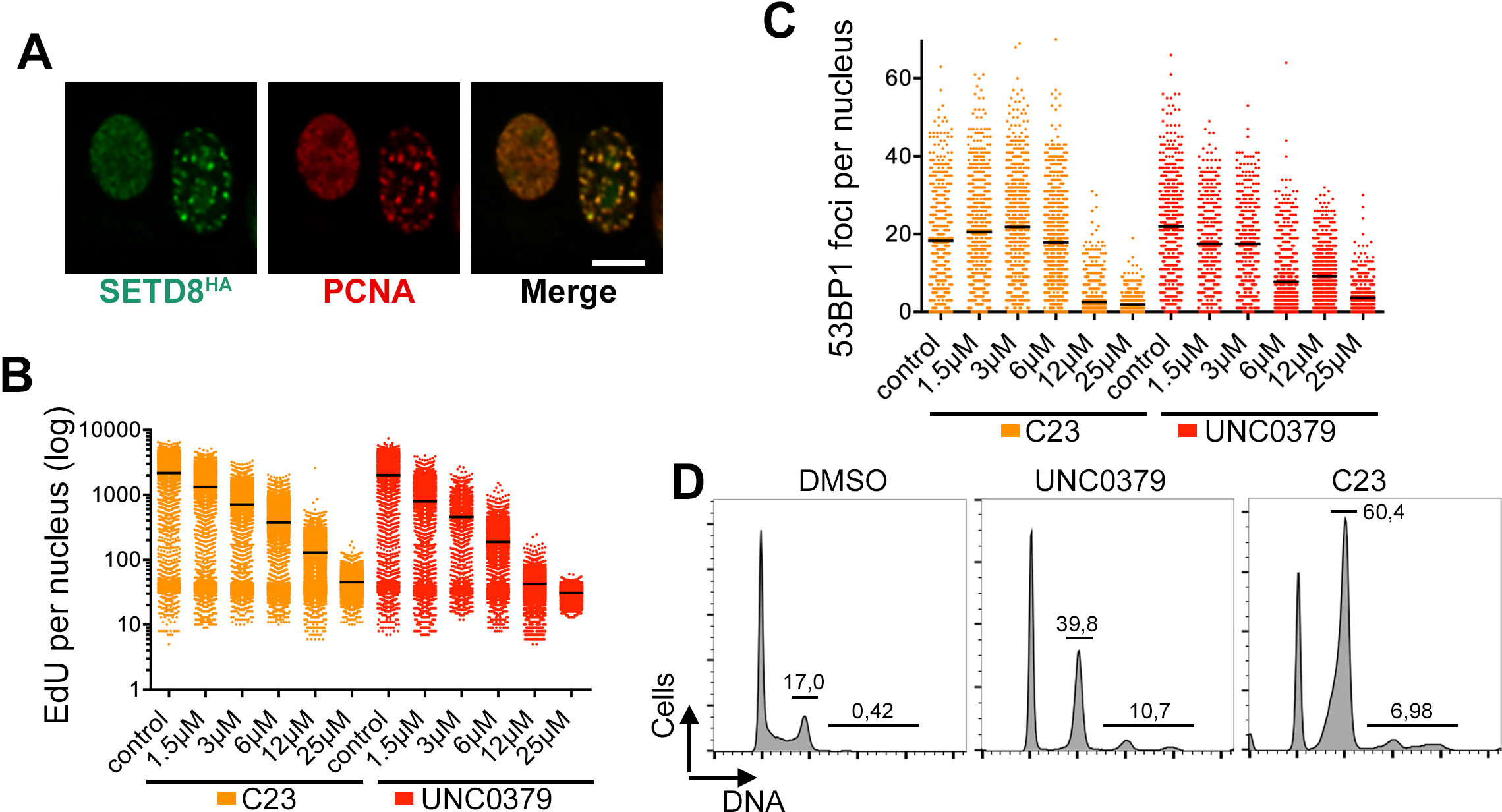
C23 phenocopies the impact of the SETD8 inhibitor UCN0379 in DNA replication. (**A**) Immunofluorescence of an HA-tagged SETD8 (SETD8^HA^, green) and PCNA (red) in U2OS cells, illustrating the localization of SETD8 to DNA replication foci. Scale bar (white) indicates 2,5 μm. (**B**) HTM-dependent quantification of the mean signal of EdU per nucleus, in U2OS cells exposed to increasing concentrations of C23 or UNC0379 for 45 min. (**C**) HTM-dependent quantification of number of 53BP1 foci per nucleus, in U2OS cells exposed to 10 Gy of IR and grown in the presence or absence of increasing concentrations of C23 or UNC0379 for 45 min. (**D**) Representative flow cytometry-based analysis of the DNA content in U2OS cells exposed for 48 h to 10 μM of UNC0379 or C23. Numbers indicate the fraction of cells in G2 and the polyploid fraction.

Given that the effects of C23 are similar to those of UNC0379, which is a substrate-competitive SETD8 inhibitor, we hypothesized that C23 could be preventing the interaction of SETD8 with chromatin. In agreement with this, fractionation experiments revealed that C23 and UNC0379 led to a dose-dependent reduction in chromatin-bound SETD8 (**Fig. 4A**), an effect that could also be seen by HTM-dependent quantification of chromatin-bound SETD8^GFP^ (**Fig. S2B**). Moreover, both compounds decreased H4K20me1 levels to a similar extent as assessed by WB (**Fig. 4B**). To specifically analyze *de novo* monomethylation of H4, U2OS were first arrested in S-phase by treatment with 1mM hydroxyurea (HU) for 24 hours, which reduces basal H4K20me1 levels, and we monitored the rescue of H4K20 methylation 6 hours after releasing the cells from the HU arrest. Using this pipeline, HTM-dependent quantification of H4K20me1 levels confirmed a dose-response effect of C23 in inhibiting *de novo* H4K20 methylation (**Fig. 4C,D**). Finally, we used the published structure of a derivative of UNC0379 (MS2177) bound to SETD8 [18], and showed that C23 or its analog C110 could potentially occupy the same pocket (**Fig. S2C**). In fact, the chemical series found in our study shares a central quinoline moiety with the UNC0379 chemical series (quinazoline in this case), highlighting the key role of this region in mediating the binding to SETD8 (**Fig. S2D**). Together, these data indicate that C23 is a novel chemical inhibitor of SETD8.

**Figure 4.**
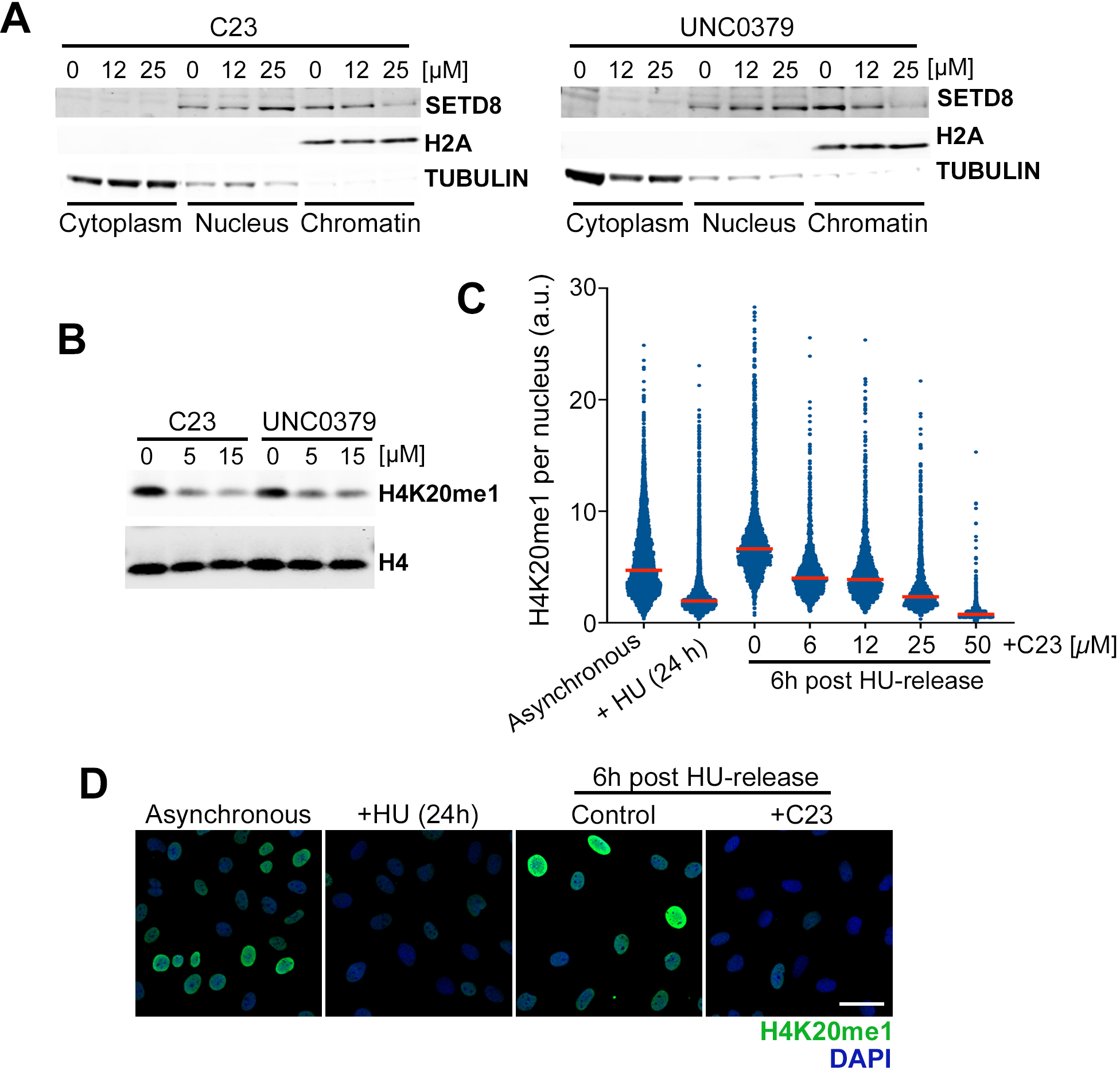
C23 is a SETD8 inhibitor. (**A**) WB illustrating the distribution of SETD8 in the cytoplasmic, nucleoplasmic and chromatin-bound fractions of U2OS cells treated with increasing concentrations of C23 or UNC0379. H2A and TUBULIN levels are shown as control of the fractionation. (**B**) WB of H4K20me1 and total H4 in U2OS cells treated with 5 and 15 μM of C23 or UNC0379 for 8 h. (**B**) HTM- dependent quantification of H4K20me1 levels per nucleus, in U2OS cells arrested with 1mM HU for 24 h, and then released in the presence of increasing concentrations of C23 for 6 h. Red lines indicate median values. (**D**) Representative images from the experiment defined in (**C**). Scale bar (white) indicates 10 μm.

### SETD8 inhibition has a particular impact on nucleoli

Early work suggested that the antitumoral effects of SETD8 inhibitors were due to the activation of p53 [9]. Consistent with this view, C23 and UNC0379 led to a similar dose-dependent increase in p53 levels in two independent neuroblastoma cell lines (NGP and SH-SY5Y) (**Fig. S3A**). However, and in agreement with other reports [20, 21], experiments in WT and p53-deficient mouse embryonic fibroblasts (MEF) revealed that SETD8 inhibitors killed cells regardless of p53 status (**Fig. S3B**). To further understand the mechanism of toxicity of SETD8 inhibitors, we performed thermal proteome profiling (TPP) in U2OS cells exposed to the C23 or its derivative C110. TPP enables the systematic assessment of proteins that change their physical properties in the presence of a chemical, thereby highlighting the cellular pathways affected by the drug [22]. These experiments were performed in duplicates, with a good correlation between both replicates for each drug (**Fig. S4A**).

Importantly, and while SETD8 peptides were only detected in the experiments done with C110, both replicates showed a significant change in SETD8 thermostability in response to the drug (**Fig. 5A**). Besides SETD8, pathway analyses using Enrichr [23] revealed an enrichment of factors related to the nucleolus and rRNA processing among those most significantly altered by C23 and C110 (**Fig. 5B**). An example is upstream binding transcription factor (UBF), a key regulatory factor for rRNA transcription, and which was thermostabilized by C23 and C110 in all tested conditions, indicating that the drugs had a particular effect on this factor (**Fig. S4B**). Consistent with TPP data, HTM analyses revealed a significant depletion of UBF in U2OS cells treated with C23, C110 or UNC0379 (**Fig. 5C, D**). Similar effects were observed in other nucleolar factors such as nucleophosmin (NPM) and fibrillarin (FBL) (**Fig. 5C**). Transmission electron microscopy (TEM) analyses confirmed numerous ultrastructural alterations of nucleoli in cells treated with C23 or UNC0379, including severe segregation of nucleolar components, a feature generally associated with a reduced rate of rRNA synthesis, and formation of both large fibrillar centers and intranucleolar vacuoles (**Fig. 5E** and **S4C**).

**Figure 5.**
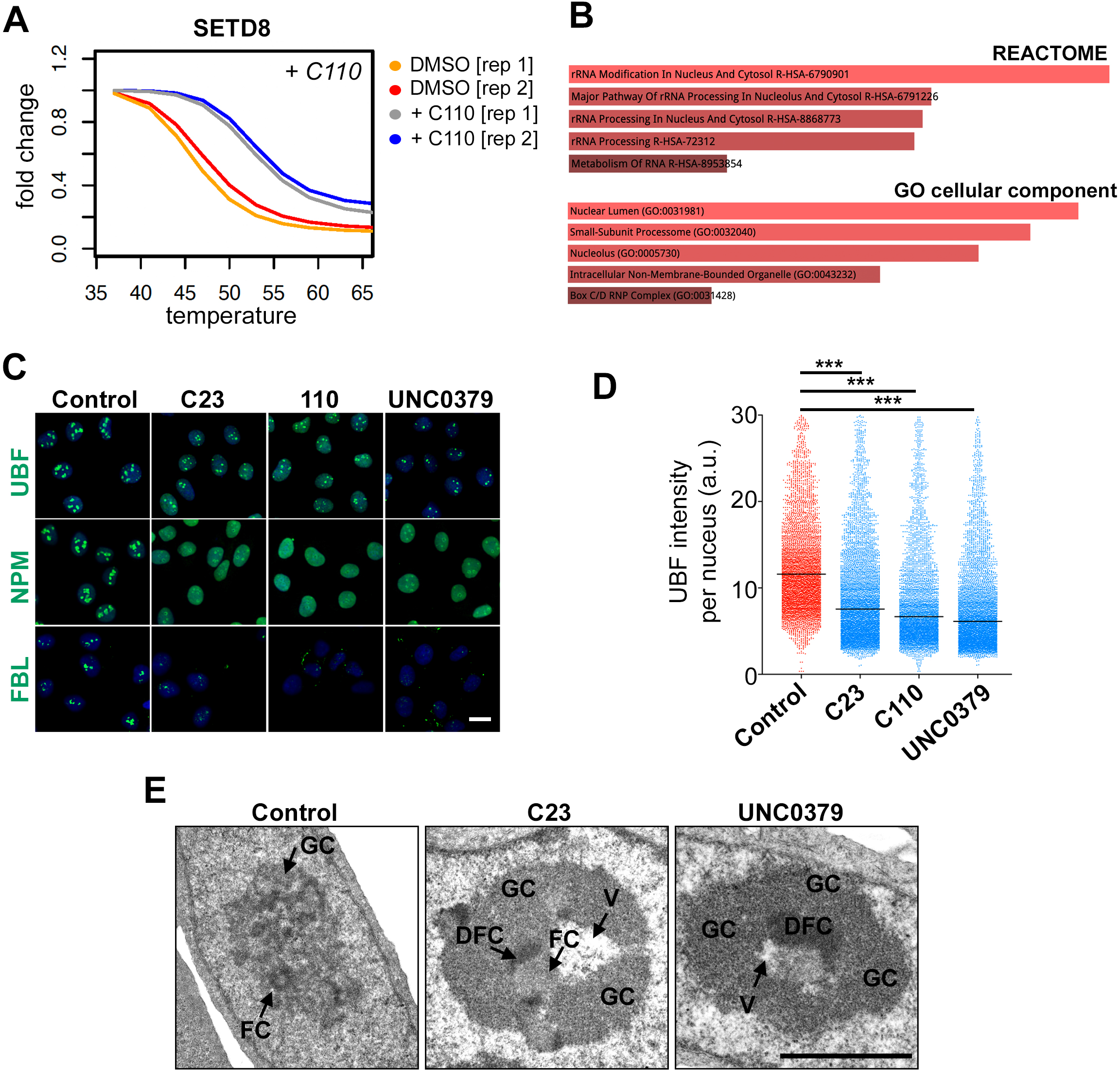
C23 has a preferential impact on nucleolar factors. (**A**) Impact of increasing the temperature on SETD8 levels as assessed by TPP in U2OS cells treated with C110 (20 μM) for 25 min. Data from 2 biological replicates is shown. (**B**) Enrichr analysis from the list of factors that presented an altered thermostability in the presence of C23. The most significant classes from “REACTOME” pathways and “GO cellular component” are shown, indicating a distinct effect of C23 on nucleoli and rRNA metabolism. (**C**) Immunofluorescence images from UBF, NPM and FBL (green) in U2OS cells exposed for 45 min to C23, C110 or UNC0379 at 10 μM. DAPI (blue) was used to stain DNA. Scale bar (white) indicates 5 μm. (**D)** HTM-dependent quantification of UBF1 levels per nucleus, from the experiment defined in (**C**). Black lines indicate median values. (**E**) Representative TEM images of U2OS cells treated with C23 or UNC0379 (25 μM) for 45 min. Control cells exhibit typical reticulated nucleoli with several small fibrillar centers (FC), surrounded by a shell of dense fibrillar component (DFC), and small irregular masses of granular component (GC). Treatment with the drug (C23 or UNC0379) induced the alteration of nucleolar integrity, including the loss of compartmentalization (nucleolar segregation), together with the formation of an enlarged fibrillar centers (FC) and accumulation of large masses of the granular component (GC). The formation of intranucleolar vacuoles (V) was also a recurrent observation in the presence of either drug. Scale bar (black) indicates 2 μM. Additional examples are provided in **Fig. S4C**. ***P < 0.001; t-test.

### SETD8 suppresses nucleolar stress

Several evidences suggest that these effects of SETD8 inhibitors on nucleoli are on target. First, proteomic analyses of SETD8 interactors reported an abundance of nucleolar proteins and SETD8 depletion impaired rRNA processing [24]. In agreement with this, immunofluorescence analyses of U2OS cells identified cells with nucleolar accumulation of SETD8^HA^ which, together with UBF, was lost upon treatment with C23 (**Fig. 6A**). To address whether genetic targeting of SETD8 had an impact on nucleolar function we used mouse embryonic stem cells (mESC) where *Setd8* can be deleted upon treatment with 4- hydroxytamoxifen (4-OHT) (*Setd8^lox/lox^)* [19]. WB analyses confirmed the loss of SETD8 expression in *Setd8^lox/-^* cells upon 4-OHT treatment (**Fig. 6B**). In agreement with inhibitor data, SETD8 deletion had a particular impact on nucleoli, as evidenced by a significant reduction of the nucleolar UBF signal as well as in rRNA transcription, monitored with a short pulse of 5-Ethynyl-uridine (EU) (**Fig. 6C-F**). Moreover, TEM analysis identified numerous nucleolar abnormalities in SETD8-deficient cells, similar to the ones observed with SETD8 inhibitors, such as segregation of nucleolar components, presence of giant fibrillar centers, formation of intranucleolar vacuoles and nucleolar fragmentation (**Fig. 6G** and **Fig. S4D**). Together, these experiments indicate that targeting SETD8 generates nucleolar stress in mammalian cells.

**Figure 6.**
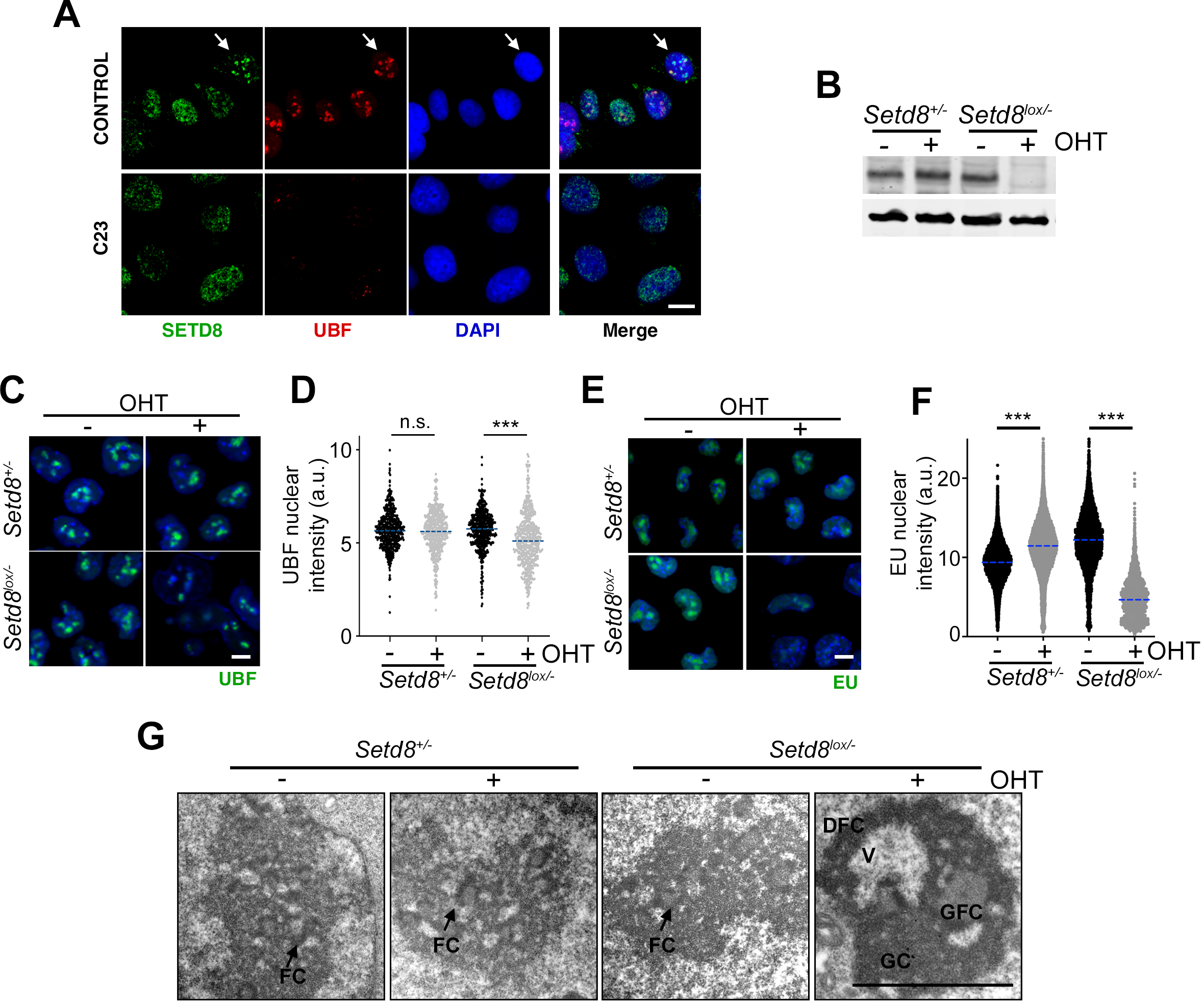
SETD8 deletion impairs nucleolar structure and function. (**A**) Representative images of the colocalization of SETD8^HA^ (green) and UBF (red) in U2OS cells, which was lost upon treatment with C23. Scale bar (white) indicates 2.5 μM. DAPI (blue) was used to stain DNA. (**B**) WB illustrating the loss of SETD8 expression in *Setd8^lox/-^* mESC 24 h after exposure to 4-OHT (1 μM). Targeted deletion was mediated by the expression of a tamoxifen inducible Cre recombinase (CreER). β-ACTIN levels are shown as a loading control. (**C**) Immunofluorescence images from UBF (green) in *Setd8^+/-^*and *Setd8^lox/-^* mESC 24 h after exposure to 4-OHT (1 μM). DAPI (blue) was used to stain DNA. Scale bar (white) indicates 2.5 μM. (**D**) HTM-dependent quantification of UBF nuclear levels, from the experiment defined in (**C**). Dashed lines indicate median values. (**E**) Immunofluorescence images of EU levels (green) in *Setd8^+/-^* and *Setd8^lox/-^*mESC 24 h after exposure to 4-OHT (1 μM). EU was added for the last 30 min before fixation. DAPI (blue) was used to stain DNA. Scale bar (white) indicates 2.5 μM. (**F**) HTM-dependent quantification of EU levels per nucleus, from the experiment defined in (**E**). Dashed lines indicate median values. (**F**) Representative TEM images from *Setd8^+/-^*and *Setd8^lox/-^* mESC 24 h after exposure to 4-OHT (1 μM). Whereas cells that preserved SETD8 expression lacked nucleolar alterations with several small fibrillar centers (FC), depletion of SETD8 induced severe nucleolar abnormalities, including prominent nucleolar segregation, appearance of both giant fibrillar centers (GFC) and intranucleolar vacuoles (V) and nucleolar fragmentation, as seen with SETD8 inhibitors. Scale bar (white) indicates 2 μm. Additional examples are provided in **Fig. S4D**. n.s.: non-significant; ***P < 0.001; t-test.

### The toxicity of SETD8 inhibitors correlates with nucleolar activity

Finally, we aimed to identify mutations that modulate the sensitivity to SETD8 inhibitors. To do so, we conducted a genome-wide genetic screen using CRISPR- Cas9 in KBM7 cells treated with C23. We chose KBM7 cells, as this is a widely used model for genetic screens with which we have experience [25]. While the screening did not find a single factor that provided a profound resistance to C23, there was a significant enrichment of factors related to nucleolar function among those that decreased the sensitivity to the compound (**Fig. 7A, B**). Based on these results, and since SETD8 inhibitors generate nucleolar stress, we hypothesized that mutations that increase nucleolar activity and ribosome biogenesis rates should increase the sensitivity to C23. In fact, one of the few conditions that has been shown to sensitize to SETD8 inhibition is MYC overexpression [9], and MYC is one of the key factors that stimulates ribosome biogenesis [26]. Consistent with this study, C23 showed increase toxicity in Ba/F3 murine lymphoma precursor cells exposed to 4-OHT, which promotes the nuclear translocation of a fusion between MYC and a fragment of the estrogen receptor (BaF^MycER^) (**Fig. 7C**). Conversely, RNA interference-mediated depletion of MYC reduced C23 toxicity (**Fig. 7D**). Besides MYC, mTOR is the other main pathway that regulates nucleolar activity and ribosome biogenesis [27]. In this regard, C23 toxicity was increased in MEF lacking tuberous sclerosis complex 2 (TSC2), where the mTOR pathway is hyperactive (**Fig. 7E**). Moreover, the increased sensitivity of C23 upon MYC expression was alleviated by the mTOR inhibitor Rapamycin in two independent murine lymphoma cell lines (**Fig. S5**). Together, these experiments indicate that ribosome biogenesis rates are a major determinant of the toxicity of SETD8 inhibitors and suggest that, in addition to MYC, mTOR activity can constitute a potential biomarker of the antitumoral effects of these compounds.

**Figure 7.**
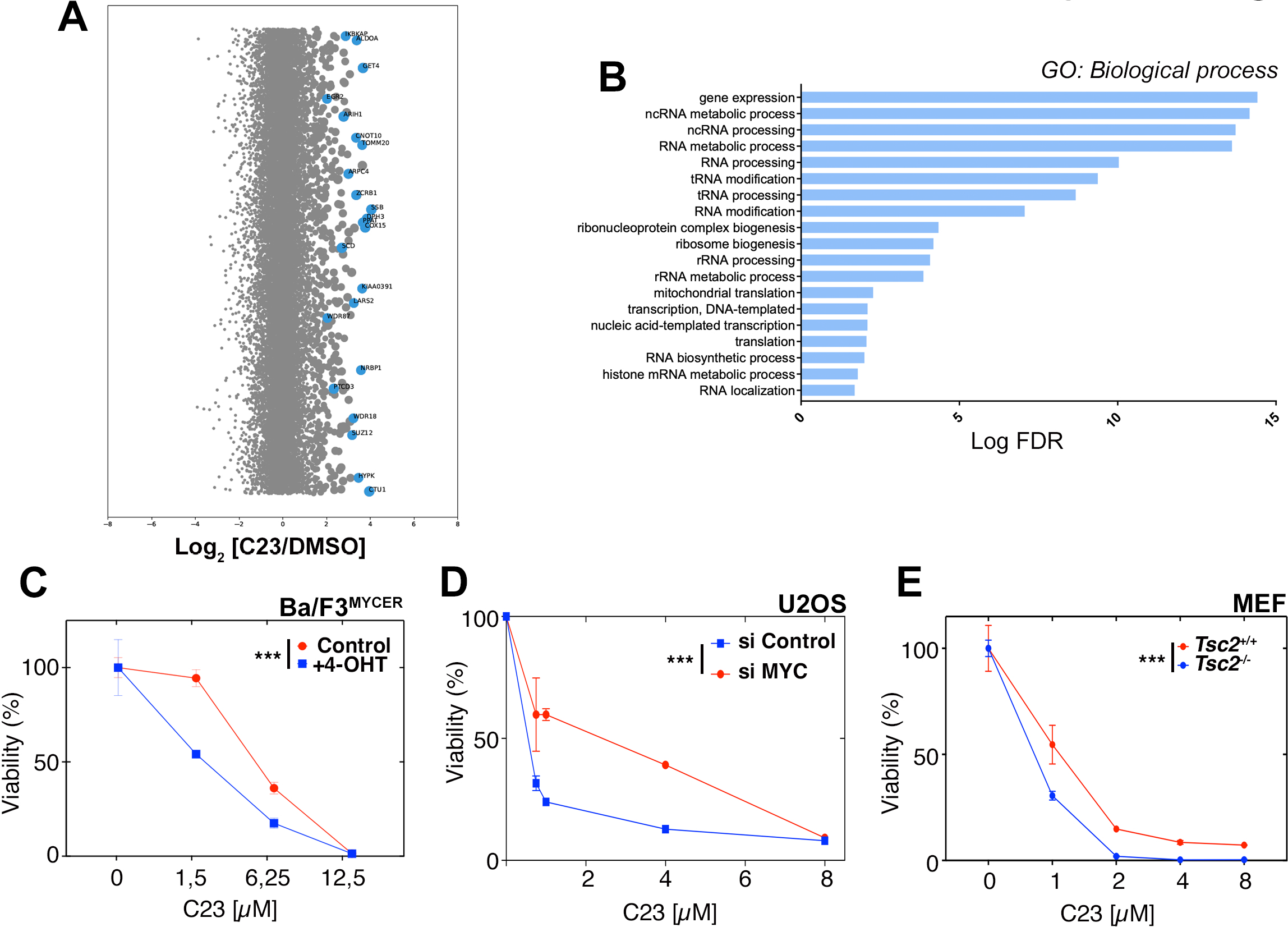
The toxicity of SETD8 inhibitors correlates with ribosome biogenesis. (**A**) sgRNA enrichment scores from a CRISPR resistance screen for C23 (10 μM) in KBM7 cells, 10 days after starting the treatment. Bubble plots illustrate the median enrichment of all sgRNAs targeting a given gene over DMSO, bubble sizes indicate significance. Significantly enriched sgRNAs (blue bubbles) were used for the enrichment analysis shown in (**B**). (**B**) Gene Ontology “Biological process” pathways that are significantly enriched among the list of significantly enriched sgRNAs selected from (**A**). Note that most pathways are related to RNA metabolism and ribosome biogenesis. (**C**) Effect of increasing doses of C23 on the viability of Ba/F3^MYCER^ cells, in the presence or absence of 4-OHT to promote the nuclear translocation of MYC^ER^. (**D**) Effect of increasing doses of C23 on the viability of U2OS cells, previously transfected with siRNAs targeting MYC or a control siRNA. (**E**) Effect of increasing doses of C23 on the viability of *Tsc2*^+/+^ and *Tsc2*^-/-^ MEF. Note that both cell lines were also p53 deficient, which is necessary to enable the growth of *Tsc2*^-/-^ MEF. ***P < 0.001; ANOVA.

## DISCUSSION

Despite intense basic research efforts done in epigenetics, this has unfortunately translated in limited treatments for human disease. This is particularly the case for inhibitors of histone methyltransferases, where only EZH2 inhibitors have progressed to clinical use [28]. Regardless of the availability of potent inhibitors, part of the limitation is the lack of biomarkers that can help to identify the type of tumor or specific patients that are most likely to respond to a given therapy. This is also the case for SETD8 inhibitors, where available compounds have poor pharmacological properties, and none have progressed to clinical trials. In this context, our work provides a novel class of SETD8 inhibitors that, to the very least, can be a useful research tool, but also helps to identify a potential biomarker of sensitivity to this therapy.

Initial work on SETD8 inhibitors suggested that their efficacy was due to p53 activation [9], yet this hypothesis has been challenged by several works [20, 21] and also refuted by our own data. Besides p53, MYC overexpression was also proposed as a determinant of SETD8 inhibitor sensitivity [11], although the reason behind this remained to be clarified. The research presented here supports a model whereby SETD8 inhibitors generate nucleolar stress, so that their toxicity is modulated by mutations or perturbations on pathways that modify nucleolar activity and ribosome biogenesis rates, such as MYC or mTOR. Of note, a recent manuscript also reported nucleolar alterations in response to UNC0379 [20], although this was limited to one compound and was not supported by genetic data. We here present evidence that the capacity to generate nucleolar stress is extensive to several SETD8 inhibitors and, more importantly, that this is an on-target effect as similar observations were made upon genetic deletion of SETD8.

In the context of this mechanism of action, mutations that increase ribosome biogenesis (e.g. MYC overexpression, or mutations that lead to mTOR hyperactivation) are expected to sensitize to SETD8 inhibition. It would be interesting to see whether mTOR-dependent signaling events that can be monitored in biopsies, such as evaluating the phosphorylation of mTOR targets like S6K or 4EBP1, can be used as biomarkers of SETD8 inhibitor sensitivity. Moreover, given that there are numerous efforts to develop novel chemotherapies that target nucleoli or rRNA transcription [29], to what extent the toxicity of these therapies is also modulated by MYC or mTOR activity emerges as an interesting possibility.

Finally, and despite of having worked on characterizing the compounds presented here, and on the potential efficacy of SETD8 inhibitors for cancer therapy, we would also want to add a word of caution on this regard. SETD8 is essential for mouse development [19], and has also been shown to be essential in multiple tissue-specific knockout models [30–35]. Data available at the cancer dependency map (DEPMAP) [36] indicate that this essentiality occurs at the cell level, and likely to be found in all cell types. The broad impact of SETD8 inhibitors in DNA replication, repair or nucleoli, supports this point. In this context, it is expected that on-target toxicity might be significant for this therapy. Having said that, the fact that an enzyme is essential does not invalidate it as a target in chemotherapy, as indicated by the success of drugs targeting key cell-cycle regulators like topoisomerases or many of the so-called targeted therapies, which target cell-essential factors. Nevertheless, the essential nature of SETD8 makes it even more important to have a good understanding of what are the determinants of sensitivity to SETD8 inhibitors, so that the use of these drugs is directed to cancer patients with mutations that make them more likely to respond. We hope that our work brings some light in this regard.

## MATERIALS AND METHODS

### Cell lines and culture

All cells were cultivated at 37°C, in a humidified air atmosphere with 5% CO2, unless otherwise specified. U2OS (ATCC), MCF-7 (ATCC), SH-SY5Y (ATCC), and NGP (DSMZ) cells were cultivated in standard high glucose Dulbecco’s Modified Eagle Medium (DMEM) supplemented with 10% foetal bovine serum (FBS), 2 mM L-glutamine and 1% penicillin/streptomycin. Culture of SH-SY5Y and NGP cells required coating of the plates with 0.1 gelatine. KBM7 cells (kind gift of Thijn Brummelkamp) were cultivated in Iscovés modified dulbeccós medium (IMDM) supplemented with 10% foetal bovine serum (FBS), 2 mM L-glutamine and 1% penicillin/streptomycin. *Setd8*^+/-^ and *Setd8*^lox/-^ mESC (kind gift of D Reinberg)[19] were grown over 0.1 gelatin with DMEM (high glucose) supplemented with 15% knockout serum replacement (Invitrogen), LIF (1000 U/ml), 0.1 mM non-essential aminoacids, 1% glutamax and 55 mM b-mercaptoethanol. The murine B-lymphoid cell lines FL5.12 (CVCL_0262) and Ba/F3 (CVCL_0161) expressing Myc^ER^ (FLMycER, BaFMycER) were a kind gift from Bruno Amati[37] and grown in regular RPMI medium (Euroclone, Pero, Italy), which includes 2 mM glutamine and 11 mM glucose, supplemented with 10% fetal bovine serum (FBS) and, for FL5.12 and Ba/F3 cells, 2 and 1 ng/ml murine interleukin 3 (PeproTech, Rocky Hill, NJ, USA), respectively. MEF lines were cultivated in DMEM supplemented with 15% FBS, and under hypoxic conditions (5% CO2, and 5% O2). p53^-/-^ MEFs were generated from mouse embryos extracted at day 14 post-coitum. MCF7^(HA)SetD8^ cells were generated by first infecting MCF7 cells with lentiviruses expressing a SETD8 cDNA tagged with HA, followed by targeting of endogenous *SETD8* using a LentiCrisprV2_Blasti vector with two sgRNA targeting human *SETD8*.

### Cell viability

For, U2OS, MEF and Ba/F3 cells, viability was quantified by HTM-dependent quantification of nuclei following DAPI staining at the end of the treatment. For experiments related to MYC overexpression and depletion, viability was quantified a luminescent system following manufactureŕs instructions (CellTiter-Glo, Promega). For Ba/F3 and FL5.12 lymphoma precursor cell lines, viability was also measured by flow cytometry based on dye (DAPI) exclusion on live cells.

### siRNA transfection

Exponentially growing cells U2OS were trypsinised and transfected in suspension with 50 nM of control siRNAs or human siRNAs targeting C-MYC (Horizon Discovery Biosciences, ON-TARGETplus siRNAs), following manufacturer’s instructions using Lipofectamine RNAiMAX (Thermo Fisher Scientific) and OPTIMEM medium (Life Technologies).

### Western Blotting

Cell pellets were recovered and washed with cold PBS, before lysis at 4°C on a shaker, using Urea buffer (50 mM Tris, pH 7.5, 8 M urea, and 1% CHAPS). Protein concentrations were quantified using the Bio-Rad Protein Assay (Bio-Rad). Approximately 20 µg of sample was mixed with NuPAGE LDS (LifeTechnologies) and 10 mM dithiothreitol (DTT) (Sigma) and incubated at 70°C for 10 minutes. The extracts were resolved in precast 4-20% gradient polyacrylamide gels (Invitrogen) and transferred using standard methods. After blocking, the membrane was incubated overnight at 4°C with the primary antibody, and 1 h at room temperature with the secondary. Fluorophore-conjugated secondary antibodies were used for detection, using the Li-Cor LCx system. A list with the antibodies, kits, plasmids and other reagents used in this study is available at **Table S2**.

### Virtual screening and molecular dynamics simulations

A set of three initial 53BP1 structures bound to a H4 peptide with the dimethyl- lysine moiety were chosen. Namely, 2 X-ray structures (PDB codes corresponding to 2ig0 and 3lgl) [38, 39] and 1 NMR-solved structure (2lvm) [13]. Before starting molecular dynamics (MD) simulations, we checked for the presence of non-standard amino acids, cofactors, and other small molecules for all structures. The presence of structural water molecules was assessed using the program CMIP [40], which predicts those water molecules located close to or within the binding site mediating potential protein-ligand interactions. Finally, independent MD simulations were performed neglecting and/or keeping those water molecules in their original positions. Once the target preparation was finished, we proceeded to solvate and neutralize with Na^+^ ions the complex, to minimize and to run unrestrained MD simulations using a combination of the latest AMBER force fields [41–43]. In order to generate an initial ensemble of conformations we extended the simulations to 50 ns. Afterwards we checked by RMSD and B-factor analysis that the targets kept their original global structure, and no big structural rearrangements took place around the binding site.

In parallel with target preparation, the organization and cleaning up of the set of compounds was undertaken for *in silico* screening. A virtual compound collection (VCC_1_) from the ZINC database [44] containing dimethylammonium was selected based on commercial availability. This subset was expanded to include different protonation, stereo and conformational states, and was finally composed of 907,987 molecules. For this enriching procedure of the VCC_1_ set we used LigPrep from Schrödinger (LigPrep version 2.8, Schrödinger, LCC; New York, NY, 2013).

In order to include receptor variability around the binding site, we extracted a set of target structures that represented dominant conformations using a clustering analysis based on a hierarchical agglomerative approach. The clustering analysis is implemented within the AMBER module *cpptraj* [45]. A RMSD-clustering was performed on the residues that belong to the binding site. The centroid of each cluster was chosen as the cluster representative structure and the most dominant was used as rigid template for docking experiments.

Docking calculations

As stated above, after the clustering analysis, the resulting structures were subjected to docking calculations after removing the peptide in the binding site.

Besides these MD-derived structures, we also used as rigid-docking receptor, the original structures. Again, the influence in docking results of structural water molecules was assessed by keeping when necessary those water molecules. The docking program used for these experiments was Glide (Glide, version 6.1, Schrödinger, LLC, New York, NY, 2013) [46–48]. After docking calculations, sorted docking results by the *Glide score* parameter were selected and the top 50 poses were visually inspected. A final subset of 25 molecules was selected from VCC1 set and tested for biological activity.

### Modeling the interaction of SETD8 with C23 and C110

We modeled the interaction of SETD8 with C23 and C110 by using as the atomic structure of human SETD8 in complex with MS2177 (PDB ID 5T5G) as a reference. For this, C23 and C110 were computational aligned with the structure of MS2177 in complex with SETD8 using the central quinoline moiety common to the three compounds and the tools provided by UCSF ChimeraX[49]. In the models, the quinoline moiety of MS2177, C23 and C110 were placed in the same position and the structure of SETD8 was left unchanged. Panels and representations of these models were performed with UCSF ChimeraX.

### CRISPR screen

250 million KBM7 cells expressing Cas9 were transduced at a 0.3 MOI with the Human CRISPR Knockout Pooled Library (Brunello, Addgene73178, two-vector system), yielding a calculated library representation of 668 cells per sgRNA (library representation = 50 million cells). For transduction, 65µl of viral supernatant was added to 5 million cells seeded in 6-well plates in 1.5 mL of IMDM and 8µg/mL of polybrene. Cells were centrifuged at 2000rpm for 1h and incubated at 37°C overnight in a final volume of 3mL IMDM. 24h after, cells were pooled and diluted in T75 flasks. Next day, pools were selected with 1µg/mL puromycin for 5 days. 50 million cells per condition were seeded after antibiotic selection at a seeding density of 500,000 cells/mL: DMSO as a control and 1μM SETD8i C23. Uninfected control cells were also seeded in parallel with the drug, to determine the resistance. Every 2 days cells were counted, and 50 million cells were seeded again in 100mL IMDM, adding fresh drug. Treatment-resistant pools were harvested and stored at -80°C on day 9 for further analyses.

To identify the sgRNAs in the control and resistant pools, DNA was extracted using a Gentra Puregene Blood Kit (Qiagen, 158445), following manufacturer’s instructions. The U6-sgRNA cassette was then amplified by PCR using the KAPA HIFI Hot Start PCR kit (Roche, KK2502) and different tagged primers required for the subsequent Illumina sequencing. The PCR product was precipitated with sodium acetate 3M in EtOH 100% at −80°C for at least 20 min, pelleted, and resuspended in water prior to purification in agarose gel. Purified samples were sent to Illumina sequencing for gRNA detection.

Raw reads were processed and analyzed following the protocol from previously described loss-of-function CRISPR screenings (Mayor-Ruiz et al., 2020). Raw read files were converted to fastq format using the convert function from bamtools (v2.5.0)36. Sequencing adapters were trimmed using cutadapt (v2.7.12) with -g CGAAACACCG and --minimum-length = 10 (genome-wide screens only). The 20bp of spacer sequence were then extracted using fastx toolkit (v0.0.14) (http://hannonlab.cshl.edu/fastx_toolkit/) and aligned to the respective sgRNA index using bowtie2 (v2.2.4)37 allowing for one mismatch in the seed sequence. Spacers were counted using the bash command ‘cut -f 3 (0)

| sort | uniq -c’ on the sorted SAM files. A count table with all conditions was then assembled, and the counts + 1 were converted to counts-per-million to normalize for sequencing depth. Log2-normalized fold changes compared to DMSO were calculated for each spacer. Statistical analysis was performed using the STARS algorithm v1.329. For this, spacers were rank-ordered based on log2 fold change and tested with the parameters --thr 10 --dir P against a null hypothesis of 10000 random permutations. Genes with q-value < 0.05 were called as hits.

### Thermal Proteome Profiling

U2OS cells were distributed into T175 flasks in 50 mL RPMI medium. Cells were incubated with C23 (50µM) and C110 (20µM) or DMSO (vehicle) for 30 min at 37°C and 5% CO2. Cells were harvested and pelleted at 300 g and RT for 3 min and washed two times with 37°C Hank’s balanced salt solution. Cells were resuspended to a density of 30*10e6 cells/mL and distributed as 80 μL aliquots into 0.2-mL PCR tubes. One of each of the compounds and vehicle containing tubes was heated in parallel for 3 min to the respective temperatures (37, 41, 44, 47, 50, 53, 56, 59, 63, 67°C), followed by a 3-min incubation time at RT. Afterwards, cells were flash-frozen in liquid nitrogen. Cells were thawed at 25°C and lysed by this freeze-thawing cycle repeated for another three times. Cell debris and precipitated proteins were removed by centrifugation at 21000 g and 4°C for 25 min. Supernatants were transferred to new tubes and protein concentrations were determined. Equal volumes of each condition that correspond to 80 μg protein in the 37°C sample were transferred to new tubes and subjected to the following digestion. First, the samples were diluted to contain 50 mM TEAB, 0.1% SDS and 5 mM TCEP. Reduction was performed at 65°C for 30 min. The samples were then cooled down to RT and alkylated with 15 mM of chloroacetamide for 30 min. The proteins were digested overnight with 1 to 40 Lys-C to protein-ratio and consecutively with Trypsin at a 1 to 25 enzyme to protein ratio. The digested peptides were labeled by 10-plex TMT-tags using 0.6 mg of the respective label for each sample. Labelling efficiency was determined by LC-MS/MS before pooling of the samples. For the sample clean-up step, a solid phase extraction (SPE strata-X-C, Phenomenex, Torrance, CA, USA) was performed and purified samples were dried in a vacuum centrifuge. An aliquot of approximately 10 µg was suspended in LC mobile phase A (3% acetonitrile in 0.1% formic acid) and 1 µg was injected on the LC-MS/MS system. Labeled peptide extracts were combined into a single sample per experiment (C23, C110, DMSO). Aliquots of approximately 250 µg peptides were pre-fractionated by means of HiRIEF [50] using gel strips with pI range of 3-10, dried using a vacuum centrifuge resulting in 72 fractions that where pooled into 32 fractions. The fractions were injected into an LC-MS/MS measurement. Online LC-MS was performed as previously described 73 using a Dionex UltiMate™ 3000 RSLCnano System coupled to a Q-Exactive-HF mass spectrometer (Thermo Fisher Scientific, Waltham, MA, USA). Each of the samples was dissolved in 20 μL solvent A and 10 μL were injected. Samples were trapped on a C18 guard-desalting column (Acclaim PepMap 100, 75μm x 2 cm, nanoViper, C18, 5 µm, 100Å, Thermo Fisher Scientific, Waltham, MA, USA), and separated on a 50 cm long C18 column (Easy spray PepMap RSLC, C18, 2 μm, 100Å, 75 μm x 50 cm, Thermo Fisher Scientific, Waltham, MA, USA). The nano capillary solvent A was 95% water, 5%DMSO, 0.1% FA; and solvent B was 5% water, 5% DMSO, 95% acetonitrile, 0.1% FA. At a constant flow of 0.25 μl min−1, a curved gradient from 3 to 8% B (in 2 min, curve of 4) and from 8% to 45% B (in 148 min, curve of 5) was used followed by a steep increase to 99% B in 2 min. FTMS master scans were performed in a mass range of 3000-1500 m/z applying a resolution of 60 000 (and mass range 300-1500 m/z), followed by data-dependent MS/MS (35 000 resolution) on the top 5 ions using higher energy collision dissociation (HCD) at 30% normalized collision energy. Precursors were isolated with a 2 m/z window and 0.5 m/z isolation offset. Automatic gain control (AGC) targets were 1e6 for MS1 and 1e5 for MS2. Maximum injection times were 100 ms for MS1 and 100 ms for MS2. The entire duty cycle lasted ∼2.5 s. Dynamic exclusion was used with 30 s duration. Precursors with unassigned charge state or charge state 1 were excluded.

The LC-MS/MS raw data was searched using Sequest/Percolator using the software platform Proteome Discoverer (PD) version 1.4 (Thermo Fisher Scientific, Waltham, MA, USA) against a UniProt (www.uniprot.org) *Homo sapiens* reference proteome database with canonical and isoforms with 42130 entries downloaded on 2016-10-24. Oxidation of Methionine was used as dynamic modification, and carbamidomethylation of cysteine, and TMT6plex on peptide N-terminus and on Lysines used as static modifications. Isobaric tag based quantitative analysis was done using the “Reporter Ions Quantifier” node in PD. The results were analyzed using R to plot melt curves and identify proteins with thermal stability changes using a workflow that was previously described[51] except that the R2 cut-off for treated curves was set to 0.5 to accommodate melt curves where compound stabilization yields an incomplete curve.

### Transmission electron microscopy

Cells were fixed with 3% glutaraldehyde in 0.12 M phosphate buffer (PB) during 30min at room temperature. Later, they were scraped, collected in Eppendorf tubes, centrifuged to obtain a pellet, washed in PB and postfixed in 2% osmium tetroxide. Cell samples were then dehydrated with acetone and embedded in ACM Durcupan (Fluka, Sigma-Aldrich) resin. Ultrathin sections (50- 70nm) were obtained using the UltraCut UC7 ultramicrotome (Leica, Microsystems, Germany), picked up on copper grids, stained with uranyl acetate and lead citrate and examined with the JEM 1011 (JEOL, Japan) electron microscope, operating at 80 kV. Micrographs were taken with a camera (Orius 1200A; Gatan, USA) using the DigitalMicrograph software package (Gatan, USA). Electron micrographs were processed using Adobe Photoshop CS6 (v.13.0.1) (Adobe Systems).

### Data availability

The mass spectrometry proteomics TPP data have been deposited to the ProteomeXchange Consortium via the PRIDE partner repository[52] with the dataset identifier PXD051199 (Username: and Password: OxCi8cm7).

### Statistics

All statistical analyses were performed using Prism software (GraphPad Software) and statistical significance was determined where the p-value was <0.05 (*), <0.01 (**), <0.001 (***) and <0.0001 (****).

## ACKNOWLEDGEMENTS

The authors want to thank Dr. Danny Reinberg for providing the *Setd8* conditional knockout mESC lines and the proteomics, genomics, and confocal microscopy units of the CNIO for their technical support in this study. O.F-C. is supported by grants from the Spanish Ministry of Science, Innovation and Universities (PID2021-128722OB-I00, co-financed with European FEDER funds), the European Research Council (ERC-2020-PoC; 963433) the Spanish Association Against Cancer (AECC; PROYE20101FERN) and La Caixa Foundation (HR22- 00890).

## AUTHOR CONTRIBUTIONS

M.M. contributed to most experiments and data analyses, preparation of the figures and to the writing of the manuscript. G.L.-P. performed the CRISPR screen and contributed to multiple experiments. R.S., I.F., L.D. and M.O. contributed to the virtual screen. M.S. and O.L. contributed to modeling experiments. E.F. performed analyses related to nucleolar function. C.A. contributed to CSR and proteomic analyses. A.G.-S. helped with viability and molecular biology experiments. S.M., C.B.-A. and J.P. contributed to organic chemistry. M.E.A. provided technical help. B.S.-L. and R.J. contributed with the TPP experiments and analyses. M.L. helped with TEM experiments and image analyses. O.F-C. supervised the study and wrote the MS.

