## Supplementary material for "SETD8 inhibition targets cancer cells with increased rates of ribosome biogenesis": Figures S1-5; Tables S1-2

### *Murga et al* **FigS1**

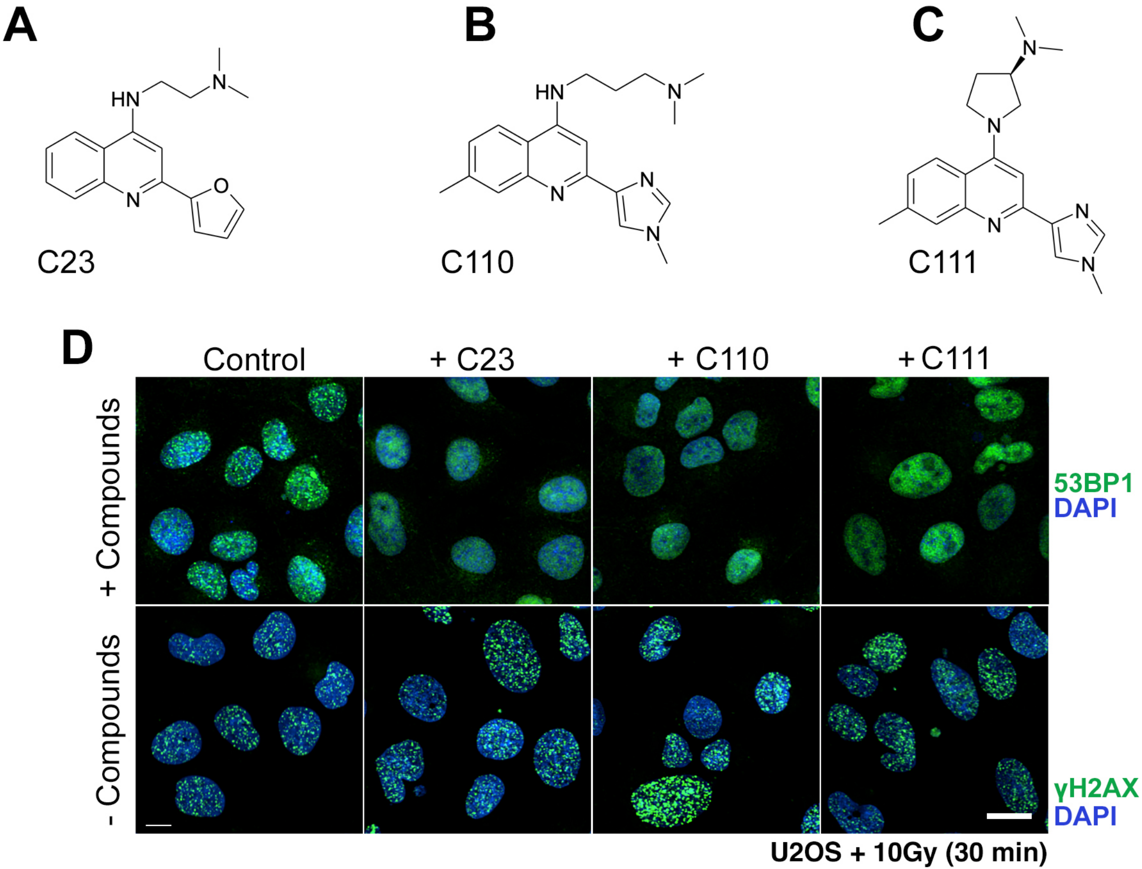

**Figure S1. C23 analogs and their effect on 53BP1 foci formation.** (**A-C**) Chemical structures of C23 and 2 analogs that are also efficient in inhibiting 53BP1 foci formation. (**D**) Immunofluorescence of 53BP1 (green) in U2OS cells, treated or not with C23, C110 or C111 (10 μM), 45 min after exposure to 10Gy of IR. Scale bar (white) indicates 5 μm.

*Murga et al* **FigS2**

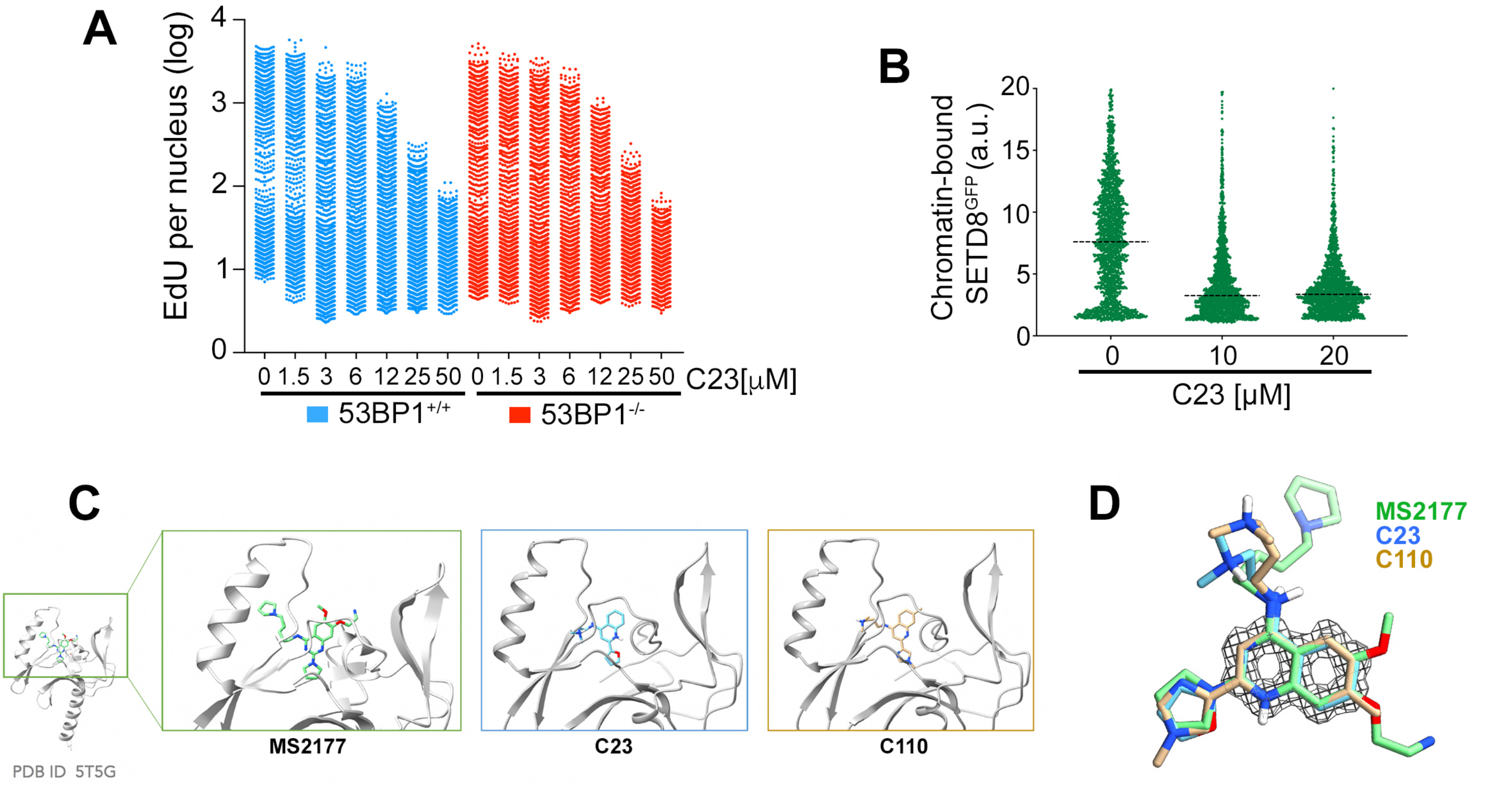

**Figure S2. 53BP1-independent functions of C23 relate to SETD8 inhibition.** (**A**) HTM-dependent quantification of EdU levels per nucleus, in *53BP1*$^{+/+}$ and *53BP1*$^{-/-}$ MEF treated with increasing concentrations of C23 for 45 min. (**B**) HTM-dependent quantification of the nuclear levels of chromatin-bound SETD8$^{GFP}$ in U2OS cells exposed for 45 min to increasing concentrations of C23. A detergent-based protocol that extracts the nuclear-soluble fraction of proteins that are not bound to chromatin was applied previous to image acquisition. (**C**) Docking of C23, C110 on the catalytic domain of SETD8, using as a reference the previously published structure of MS2177-bound SETD8 (PDB 5T5G). (**D**) Alignment of the chemical structures of MS2177, C23 and C110.

*Murga et al* **FigS3**

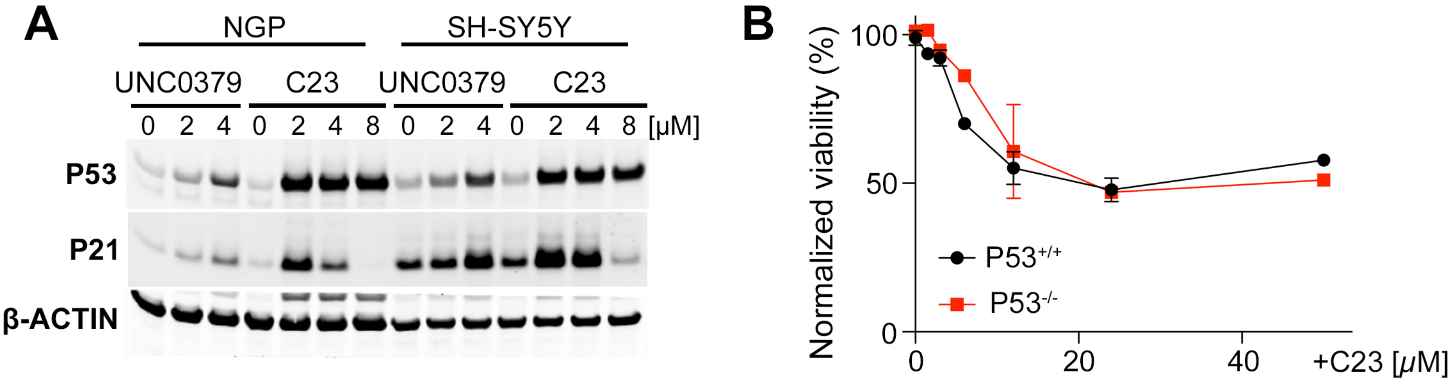

**Figure S3. C23 toxicity is p53-independent SETD8 inhibition.** (**A**) WB illustrating the levels of P53 and P21 in independent neuroblastoma cell lines treated with increasing concentrations of C23 for 8 h. β-ACTIN levels are shown as a loading control. (**B**) Effect of increasing concentrations of C23 on the viability of $P53^{+/+}$ and $P53^{-/-}$ MEF.

*Murga et al* **FigS4**

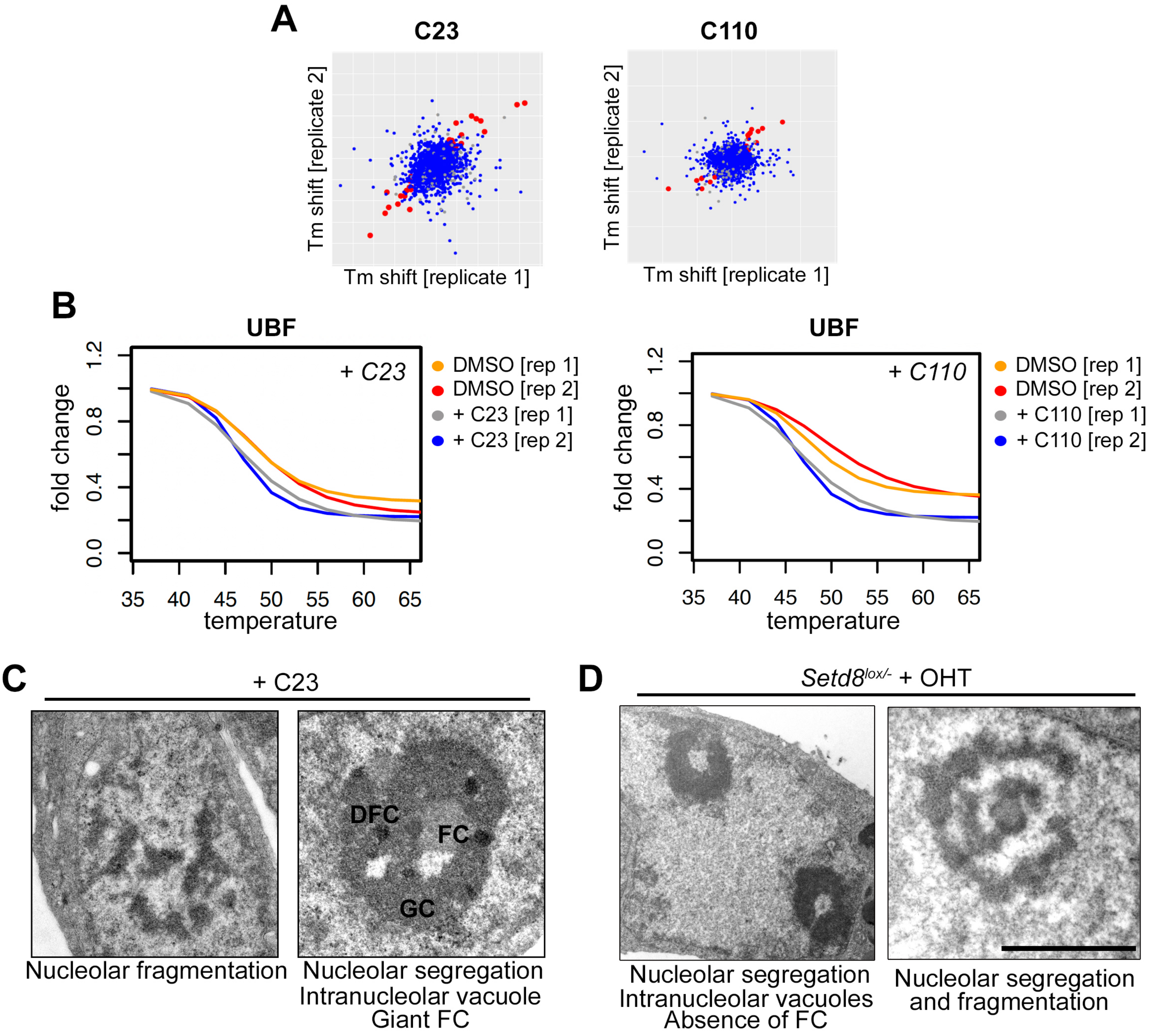

**Figure S4. Targeting SETD8 generates nucleolar stress.** (**A**) Overall correlation of melting temperature (Tm) shifts between the two biological replicates of the TPP experiment. (**B**) Impact of increasing the temperature on UBF levels as assessed by TPP in U2OS cells treated with C23 or C110 for 25 min. Data from 2 biological replicates is shown. (**C, D**) Representative TEM images from U2OS cells treated with C23 (25 μM) for 45 min (**C**) or *Setd8*$^{+/-}$ and *Setd8*$^{lox/-}$ mESC 24 h after exposure to 4-OHT (1 μM). Scale bar represents 2 μM.

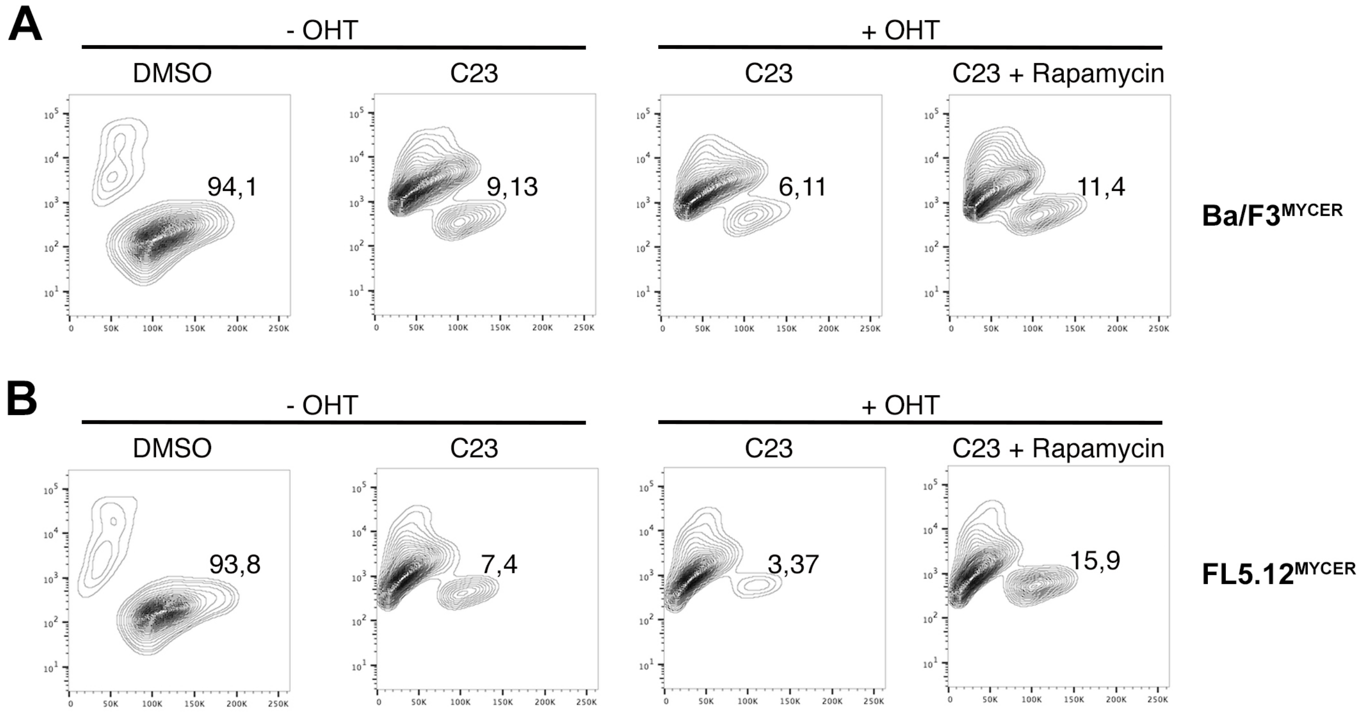
A
- OHT
+ OHT
DMSO
C23
C23
C23 + Rapamycin
94,1
9,13
6,11
11,4
Ba/F3MYCER
B
- OHT
+ OHT
DMSO
C23
C23
C23 + Rapamycin
93,8
7,4
3,37
15,9
FL5.12MYCER

**Figure S5. C23 toxicity correlates with MYC and mTOR activity.** (**A, B**) Flow cytometry profiles (*Y*-axis: DAPI; *X*-axis: FSC) of Ba/F3$^{MYCER}$ and FL5.12$^{MYCER}$ cells treated with OHT (1 mM, 24 h) in the presence or absence of C23 (10 μM) or Rapamycin (50 nM). The percentage of viable cells is indicated in each profile.

| TABLE S1. List of original hits selected from the virtual screen. | | |
|---|---|---|
| COMPOUND | NUMBER | NAME |
| AW00847 | 1 | 4-(4-fluorobenzyl)-N-phenyl-1,4-diazepane-1-carboxamide |
| RH01524 | 2 | N-((4,4-dimehtyl-2,6-dioxocyclohexylidene)methyl)aspartic acid |
| SEW05754 | 3 | 3-{[2-(4-methylpiperazino)phenyl]methylene}-1H-indol-2-one |
| RDR02556 | 4 | N1-(2,6-dimethylphenyl)-3-piperidinopropanamide |
| KS-5103 | 5 | (-)-N-(((S)-1-Ethyl-2-pyrrolidinyl)methyl)-5-sulfamoyl-o-anisamide |
| E101 | 6 | S-(–)-3-Chloro-5-ethyl-N-[(1-ethyl-2-pyrrolidinyl)methyl]-6-hydroxy-2-methoxybenzamide hydrochlo |
| M3189 | 7 | N-(2,6-Dimethylphenyl)-1-methyl-2-piperidinecarboxamide hydrochloride |
| STK134574 | 8 | 1-(diethylamino)-3-[(1,1-dioxido-1,2-benzothiazol-3-yl)amino]propan-2-ol |
| STL059340 | 9 | 3-hydroxy-N,N,3-trimethyl-4-phenylbutan-1-aminium |
| STK538232 | 10 | 1-azabicyclo[2.2.2]oct-3-yl(dithiophen-2-yl)methanol |
| STL356215 | 11 | N-(3-chlorophenyl)-2-(pyrrolidin-1-yl)acetamide |
| STK379741 | 12 | N-{4-[methyl(1-methylpiperidin-4-yl)sulfamoyl]phenyl}acetamide |
| STK887751 | 13 | 8-[(dimethylamino)methyl]-7-hydroxy-3-(1-methyl-1H-benzimidazol-2-yl)-2H-chromen-2-one |
| STL059340 | 14 | 3-hydroxy-N,N,3-trimethyl-4-phenylbutan-1-aminium |
| STK054778 | 15 | 2-{3-[3-(dimethylamino)propyl]-2-imino-2,3-dihydro-1H-benzimidazol-1-yl}-1-phenylethanol |
| STK540953 | 16 | 8-{[2-(dimethylamino)ethyl]amino}-7-ethyl-1,3-dimethyl-3,7-dihydro-1H-purine-2,6-dione |
| STL169956 | 17 | N-(4-methoxyphenyl)-4-methylpiperazine-1-carbothioamide |
| AW01002 | 18 | N-[2-(dimethylamino)ethyl]-5-(2-methyl-1,3-thiazol-4-yl)thiophene-2-sulfonamide |
| AG-205/15424896 | 19 | N-[3-(dimethylsulfamoyl)phenyl]-4-oxo-3-phenyl-2-thioxo-1H-quinazoline-7-carboxamide |
| AK-968/36977036 | 20 | (2R)-3-hydroxy-2-[(2-isopropyl-4-methyl-6-oxo-1H-pyrimidine-5-carbonyl)amino]propanoic |
| PH004881lALDRICH | 21 | 4-(dimethylamino)-2,2-diphenylpentanamide |
| S561274 | 22 | 4-amino-alpha-diethylamino-O-cresol |
| SML2187 | 23 | N2-[2-(2-Furanyl)-4-quinolinyl]-N1,N1-dimethyl-1,2-ethanediamine |
| CDS020251 | 24 | 2-[(4-Methylpiperazin-1-yl)carbonyl]aniline |

**TABLE S2. List of reagents used in this study.**

| REAGENT or RESOURCE | SOURCE | IDENTIFIER |
|---|---|---|
| Antibodies | | |
| Rabbit anti-53BP1 | Novus | Cat#100-304A2 |
| Mouse anti-H2AX | EMD Millipore | Cat#05-636 |
| Mouse anti-p53 | Santa Cruz | Cat#sc-6243 |
| Mouse Anti-KAP | BD Pharmingen | Cat#610680 |
| Rabbit anti-FBL | Cell Signaling | Cat#2639 |
| Mouse anti-UBF | Santa Cruz | Cat#sc-13125 |
| Rabbit anti-NCL | Abcam | Cat#bb22758 |
| Rabbit anti-pKAP | Bethyl Laboratories | Cat#A300-767A |
| Mouse anti-actin | Sigma | Cat#A5441 |
| Mouse anti-CHK2 | Upstate | Cat#05-649 |
| Rabbit anti-PCNA | Merck | Cat# 07-2162 |
| Rabbit anti SETD8 | Sigma | Cat# HPA064495 |
| Mouse anti-H2A | Cell Signaling | Cat#3636 |
| Mouse anti-Tubulin | Sigma | Cat# T9026 |
| Rabbit anti-H4K20me1 | Abcam | Cat#9051 |
| Rabbit anti-H4K20me1 | Upstate | Cat#07-440 |
| Rabbit anti-p21 | Cell Signaling | Cat#2947 |
| Anti-Mouse IgG-555 | Bethyl Laboratories | Cat#A90-516D3 |
| Anti-Mouse IgG-488 | Invitrogen | Cat#A11001 |
| Anti-Rabbit IgG-488 | Invitrogen | Cat#A21441 |
| Anti- Rabbit IgG-555 | Bethyl Laboratories | Cat#A120-201D3 |
| Anti- Rabbit IgG-647 | Invitrogen | Cat#A21443 |
| Anti-Rat IgG-488 | Invitrogen | Cat#A21470 |
| Mouse anti-IgA-RPE | Oxford Biotechnology | Cat#104009 |
| Rat anti-HA | Merck | Cat#11867423001 |
| Chemicals, peptides, and recombinant proteins | | |
| Anti-CD40 | BD Pharmingen | 553722 |
| IL-4 | Preprotec | 214-14 |
| IL-3 | Preprotec | 213-13 |
| CellTiter-Glo | Promega | G7571 |
| Rapamycin | Alpha Aesar | J62473 |
| TGF-$\beta$ | R&D Systems | 240-B |
| UNC0379 | Selleckchem | S7570 |
| HU | Sigma | H8627 |
| 4-OHT | Sigma | H7904 |
| Critical commercial assays | | |
| Click-It EU 488 Imaging kit | Thermo Fisher scientific | C10329 |
| Gentra Puregene Blood Kit | Qiagen | Cat#158445 |
| RnaiMax | Thermo Fisher scientific | Cat#13778150 |
| Lipofectamine 2000 | Invitrogen | Cat#11668027 |
| QuantSeq 3' mRNA-Seq Library Prep Kit | Lexogen | N/A |
| Deposited data | | |
| TPP proteomics data | PRIDE repository | PXD051199 |
| Experimental models: Cell lines | | |
| U2OS | ATCC | RRID:CVCL_0042 |
| MCF7 | ATCC | RRID:CVCL_0031 |
| KBM7 | Gift from T. Brummelkamp | RRID:CVCL_A426 |
| SH-SY5Y | ATCC | RRID:CVCL_0019 |

**TABLE S2. List of reagents used in this study.**

| | | |
|---|---|---|
| NGP | Gift ATCC | RRID:CVCL_2141 |
| *P53*$^{+/+}$ MEF | This paper | N/A |
| *P53*$^{-/-}$ MEF | This paper | N/A |
| Ba/F3_MycER | Gift Bruno Amati | Donati et al. 2022 |
| FL5.12_MycER | Gift Bruno Amati | Donati et al. 2022 |
| *Setd8 +/-* mESR | Gift Danny Reinberg | Oda et al. 2009 |
| *Setd8 lox/-* mESR | Gift Danny Reinberg | Oda et al. 2009 |
| Oligonucleotides | | |
| Setd8_sg1 ACGGAGCGCCATGAAGTCCG | This paper | N/A |
| Setd8_sg2 ACGGGAGGCTCTGTACGCAC | This paper | N/A |
| Recombinant DNA | | |
| pLEX305_SETD8_dTAG | Addgene | Addgene (Backbone #91798) |
| p-EF1a-CreERT2-3Xflag-T2A-eBFP2 | Addgene | Addgene (Plasmid #170186) |
| LentiCas9_Blasti | Addgene | Addgene (Plasmid 52962) |
| pLenti CRISPR.V2_ Blasti_sg1SetD8 | This paper | Addgene (Backbone #98293) |
| pLenti CRISPR.V2_Blasti_sg2SetD8 | This paper | Addgene (Backbone #98293) |
| Software and algorithms | | |
| GraphPad Prism | GraphPad Software Inc | http://www.graphpad.com/scientific-software/prism/ |
